# I err, therefore I am: Neural signature of error-monitoring predicts self-awareness in infants

**DOI:** 10.64898/2026.02.01.702903

**Authors:** Cécile Gal, Dimitris Askitis, Katarina Begus

## Abstract

Self-reflection is central to human cognition and arguably a defining feature of human consciousness, yet the mechanisms enabling our mind to reflect upon itself and to construct a sense of self are unclear. Specifically, whether metacognition implies self-awareness, is independent of it, or emerges because of it, remains a subject of debate, fuelled by reports of metacognitive-like behaviour in preverbal infants and non-human animals. Building on evidence that 12-month-old infants show neural signatures of internal error-monitoring, we follow infants longitudinally from 12 to 18 months, when infants first start exhibiting evidence of a conceptual self-representation. Using a gaze-contingent match-to-sample paradigm with concurrent eye-tracking and EEG, we confirm that infants exhibit a neural signature of internal error-monitoring (error-related negativity, ERN) after making an incorrect decision (saccadic eye-movement) and before receiving visual feedback. Critically, we find that ERN at 12 months developmentally predicts which of the infants will evidence self-representation (mirror self-recognition) six months later. Infants’ self-recognition was predicted specifically by internal error-monitoring, and not by external feedback-monitoring (FRN), nor by infants’ age or general development. Moreover, we show that only infants who later self-recognised adjusted their behaviour post-error, by slowing their decisions and increasing visual exploration. Thus, internal error-detection, if available, can be utilised for online adjustments in information-seeking and decision-making as early as 12 months of age. These converging neural and behavioural findings support the idea that, rather than presupposing a self, early metacognition may be functionally involved in building conceptual self-awareness in humans.

## Introduction

Behaviours and neural indices consistent with the presence of metacognitive capacities have been demonstrated in preverbal human infants [1,2] as well as several animal species [3]. These findings continue to fuel theoretical debates regarding the representational and self-reflective capacities that can be assumed to underlie such behaviours and to be supported by such neural processes [4–6]. Does seeking information under uncertainty imply an awareness of the uncertainty, and thus an understanding of the mind [7]? By ascribing metacognition to human infants, primates, even birds, do we tacitly accept that these individuals are therefore self-aware [5]? Or can cognitive processes be monitored, and behaviours triggered, by neural representations of ignorance, uncertainty, or error, which can guide behaviour without these representations being conceptual nor themselves reaching or contributing to self-awareness [8]? Here, we approach this puzzle by studying human infants longitudinally and investigating whether the capacities to monitor and control own cognition and behaviour, at 12 months, are developmentally linked to the emergence of self-representation in the same infants, 6 months later. Such a predictive relationship would suggest that, rather than presupposing self-awareness, metacognition may be functionally involved in building infants’ emerging conceptual self-representation.

In developmental research, the hallmark of having acquired a conceptual representation of the self is indicated by infants’ success on the mirror self-recognition (MSR) test. Self-representation is attributed to a child if, when placed in front of a mirror while bearing a mark previously placed on their face unbeknownst to them, the child reaches for the mark on their own body [9]. Although some have questioned the relevance of MSR for conceptual self-knowledge [e.g., 10], the success on the MSR has been associated with a host of other co-emerging behaviours indicative of self-representation, such as infants using personal pronouns [11], exhibiting self-conscious emotions like embarrassment [12], and showing a self-reference memory bias [13]. Combined, the evidence therefore suggests that success on the MSR is a robust and meaningful proxy for the presence of a conceptual self-representation in infants. We capitalise on the fact that, in western cultures, approximately 50% of infants exhibit such evidence of self-representation by 18 months of age [14]. Harnessing this variability, we investigate whether earlier mechanisms of cognitive self-monitoring predict which infants will evidence conceptual self-awareness amongst age-matched 18-month-olds.

Our primary measure of infants’ cognitive self-monitoring is error-related negativity (ERN), a well-established electroencephalography (EEG) signature of internal error monitoring in adults. The ERN is a negative deflection in frontocentral brain activity originating from the anterior cingulate cortex and arising after an individual commits an error [6]. It is believed to reflect a second-order, post-decisional mechanism, signalling a discrepancy between the ongoing (or just-performed) action and the intended action [15]. Importantly, this comparison is internal and does not rely on external feedback [15]. Moreover, ERN occurs only when participants are conscious of the action they ought to perform, and absent if stimuli are subliminal, or if subjects are ignorant of the correct option [6]. In infants, one prior study has demonstrated the presence of an ERN in 12-month-olds, when infants made incorrect predictive eye-movements [16]. Like in adults, the infant ERN was not found when the stimuli were presented subliminally and occurred prior to infants receiving external feedback. Here, we measure the ERN as an index of early self-monitoring (error-monitoring) capacities in 12-month-old infants and investigate its predictive value for later emerging conceptual self-representation in the same infants, 6 months later.

## Methods

### Participants

124 participants were recruited from the database of volunteer families at our research centre. They were aged 12 months at visit 1 (average age 12 months and 3 days, S.D. 8 days, 60 females) and came back six months later for visit 2 (114 participants came back, average age 17 months and 24 days, S.D. 12 days, 56 females). 58 infants (28 females) contributed enough data to be included in the whole-group analysis on visit 1, and 53 infants could be included in the analysis for both visits (4 infants did not attend the second visit, and 1 contributed inconclusive data for the mirror-self-recognition task). Families volunteered for a large multi-task longitudinal project (only relevant tasks mentioned here) and received gifts but no monetary compensation for participation. The participants were typically developing, full-term infants (35 gestational weeks or more). Their caregivers gave informed consent before any data collection took place, and the procedures were approved by the university’s Ethical Committee.

### Mirror-self-recognition (MSR) test (18 months)

*Procedure and data processing*. The procedure (based on [17,18]) comprised 4 phases: 1) the infant was placed in front of a mirror and familiarized with it; 2) the mirror was occluded with a curtain, the infant was given a toy to play with, and a mark was applied on their nose surreptitiously while they played, using nose-blowing as a pretence, after which the toy was taken away; 3) the mirror was uncovered and the child was exposed to the mirror for at least 3 looks to their own face in the mirror or until they touched their nose; 4) the experimenter pointed to the child in the mirror and said ‘who is this?’. Children’s behaviour was video recorded and coded offline for their reactions during phase 3. The data was independently coded by 2 experimenters with 90% agreements (10 out of 111 videos coded differently). Videos where experimenters disagreed were reviewed and coded together to reach an agreement (7 videos) or marked as inconclusive (3 videos). Infants were categorized as passing the MSR test if they touched their marked nose (*S*, N=33 in the included sample, 21 females) or not passing if they did not exhibit this behaviour (*NS*, N=20, 7 females).

### Match-to-sample task (12 months; see Figure 1)

#### Procedure

Infants were sat on their caregiver’s lap, who were instructed to limit their interaction with their infants during the task. To help with infant engagement, snacks could be given and videos could be played during short breaks. Infants were placed at a 60cm distance from a 17’’ screen on which the stimuli were presented using PsychToolbox for Matlab. A video was played as the infants were sat in front of the apparatus, followed by a 5-point eye-tracking calibration before the task started. The task went on until the infant completed 50 test trials or until they became too fussy to attend (crying, getting off parent’s lap, turning away from the screen for a continuous 30s). Infants’ gaze was tracked using an Eyelink1000 Plus eye-tracker at a 1,000Hz sampling rate (SR Research, Ontario, Canada). Infants wore an electroencephalographic (EEG) 128-channel HydroCel Geodesic Sensor net (Electrical Geodesics, Inc., USA) recording their scalp potentials with a vertex reference (Cz) at 1,000Hz using EGI NetStation software.

**Figure 1.**
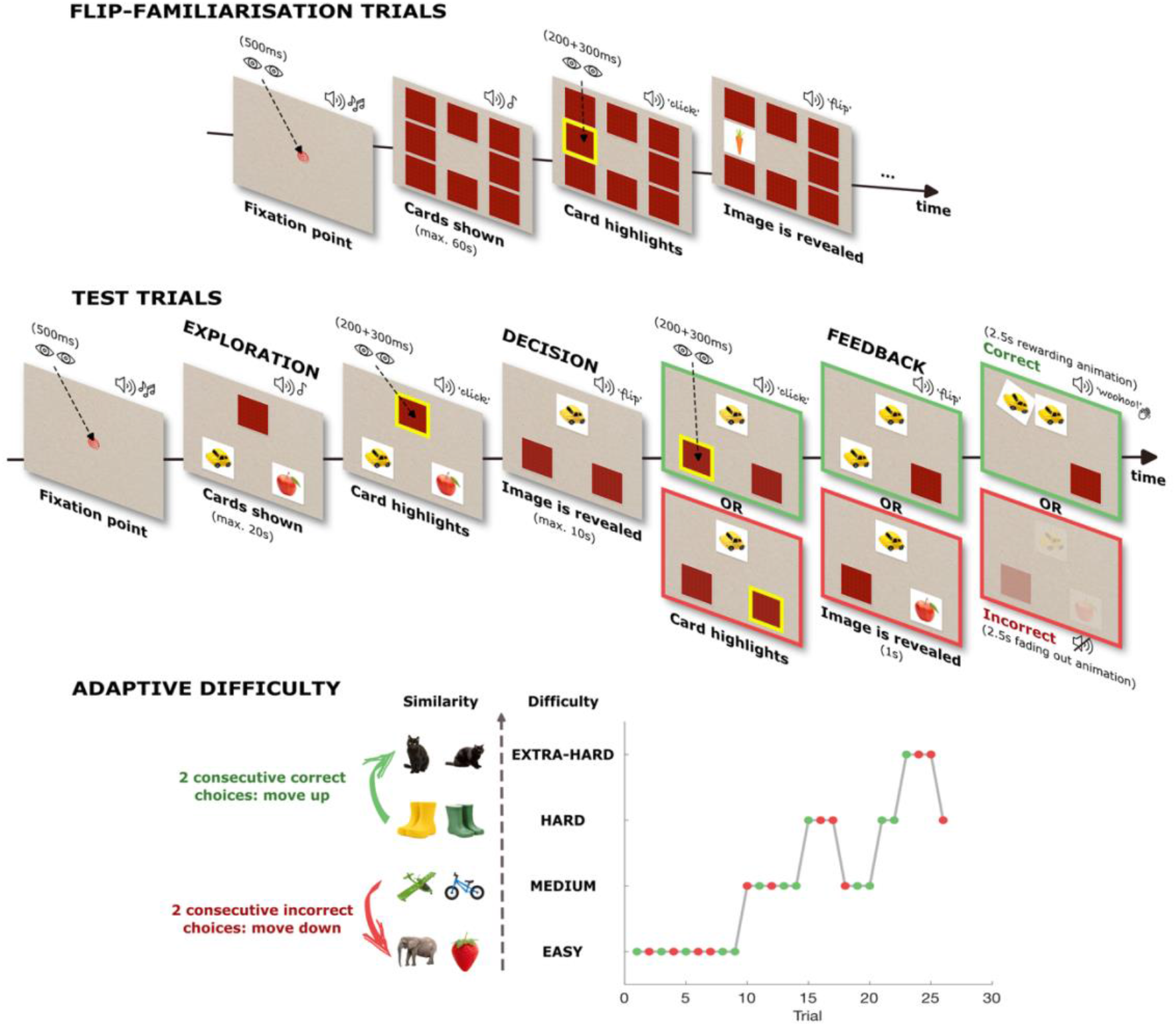
Match-to-sample task design. **Test trials:** trial sequence is represented with screenshots of different trial steps along the 3 phases (exploration, decision, feedback) described in the ‘trial design’ section; screenshots corresponding to trials where the participant triggered the correct option are highlighted in green, while screenshots highlighted in red correspond to incorrect trials. **Flip-familiarisation trials:** the trial sequence is depicted with screenshots of example trial steps. Gaze-contingent elements are represented by an arrow linking schematic eyes to the triggerable part of the screen; sounds are depicted with loudspeaker symbol where applicable. **Adaptive difficulty:** An example plot of a participants’ sequence of trials (x axis) and the corresponding difficulty level of the trials (y axis) are depicted, together with the accuracy of the participants’ response (correct trials depicted in green, incorrect in red). Example pairs of stimuli are represented along each level of difficulty. Participants started at the easiest level and followed an adaptive difficulty design based on the participants’ answers: 2 consecutive correct answers caused a one-step difficulty increase while 2 consecutive incorrect answers caused a one-step drop in difficulty.

#### Stimuli

Stimuli consisted of images of items familiar to infants, divided into different categories, subcategories and colours. Two different images were used per trial, displayed over 3 cards, such that the top card’s image was identical to one of the bottom two cards’ images. Four levels of difficulty were implemented by manipulating the similarity of the images: Easy (different categories and colour), Medium (same category, different subcategories and colour), Hard (same category and subcategory, different colours), Extra-Hard (same category, subcategory, and colour). A different set of images was used for familiarization (12 images: 3 categories, each with 4 subcategories, all different colours) and test trials (64 images: 4 categories, each with 4 subcategories, each with 4 items of same or different colours). The images (128*128 pixels) were presented on a square white background representing a play card (170*170 pixels, 205*205 pixels area of interest – AOI), and the cards were presented on a lightly textured full-screen beige background.

#### Gaze-contingency

Infants’ gaze was tracked and used to trigger the display of stimuli in real time. Before each trial, a central fixation mark was presented (50*50 pixels for a 125*15 pixels AOI). 500ms of continuous gaze detected on the central fixation triggered the trial start. During each trial, several events could be triggered if continuous gaze was detected within specified areas of interest (AOI). Bouts of up to 100ms out of the AOI were allowed without re-starting the count for optimal noise-handling. Trials were terminated if no gaze was detected on screen for 5s.

#### Test trials design

Each trial displayed 3 cards at the centre of the screen and consisted of 3 phases: 1) *Exploration phase*: The bottom two cards were presented face-up (images visible), while the top card was face-down (image not visible). Infants could explore the display for up to 20s. If gaze was detected on the face-down card, the card’s edge would highlight in yellow (after 200ms continuous gaze, accompanied with a click sound). An additional 300ms of continuous gaze on the same card would trigger the card to flip and reveal its image (with a flip sound), while simultaneously flipping the bottom two cards face-down, and starting the *Decision phase*. The first 2s of the *Exploration phase* were not gaze contingent to encourage exploration. 2) *Decision phase*: Infants could explore the display for up to 10s. If gaze was detected on one of the bottom (now face-down) cards, the card would highlight (200ms continuous gaze) and flip (after an additional 300ms continuous gaze). The time-counts to trigger the highlight and the flip were independent (infants could look elsewhere in-between highlight and reveal), but only one card could be highlighted at a time, flipping could only be triggered after highlighting, and only one of the bottom two cards could be flipped per trial. 3) *Feedback phase*: After selecting and flipping one of the bottom cards, images remained static for 1000ms (visual feedback), followed by a 2.5s valent feedback. In correct trials, a rewarding sound (applause) was played while the selected card moved up besides the top card, tilted sideways and back, then all the cards moved towards the right of the screen until they disappeared. For incorrect trials, all cards slowly faded out into the background, with no accompanying sound.

Test trials started at the easiest difficulty level, then followed an adaptive difficulty design: two consecutive correct choices lead to one level increase, and two consecutive incorrect choices lead to difficulty decreasing one level. Trials were pseudo-randomized such that each side was set to be the correct side twice every four trials, every category or subcategory appeared once before a previously shown category or subcategory could repeat, and colours and categories did not repeat from one trial to the next (or for more than two trials in a row for Easy trials where stricter control of repetitions would make the categories predictable).

#### Familiarization trials design

Prior to test trials, 6 familiarization trials were presented using a separate set of images (see ‘Stimuli’ section): 2 flip-familiarization and 4 match-familiarization trials. *Flip-familiarization trials*: 8 face-down cards were presented along the edges of the screen. Cards could be triggered as described above for test trials. Triggering the reveal of a card’s image caused any previous face-up card to flip face-down. The trial ended after all images were revealed or no card was flipped for 7 consecutive seconds or more than 30 seconds had passed. *Match-familiarization trials*: Identical to test trials (see above), except for small differences aimed at helping the infants grasp the goal of the task: 1) *Exploration phase*: identical to test trials. 2) *Decision phase*: an extra attention-getting animation was played at the start – after flipping (triggered by gaze), the top card tilted left and right while the word ‘See’ was played. 3) *Feedback phase*: selecting one card triggered the reveal of both cards (all images visible to aid rule learning), incorrect choices were accompanied by a buzzer sound, and this phase was extended such that infants were given as second chance if they first picked the incorrect choice: the matching card moved up next to the top card and back down and the trial loop restarted at the beginning of the decision phase. Match familiarization trials could be repeated throughout the task if an infant repeatedly chose cards on the same side of the screen (3 out of the last 4 test trials); test trials resumed after 4-trial match-familiarization trials or when the infant picked the side opposite to their side-bias.

### Data processing and analyses

Data was primarily analysed with Matlab (version 2024a; MathWorks, 2025), using custom scripts (available online on GitHub; see our code availability statement at the end of the article), and with JASP (version 0.19.3; JASP Team, 2025). EEG data was also pre-processed in NetStation (version 5.4.3 r29917; Electrical Geodesics, 2025) and in Matlab with the toolbox FieldTrip [version 20240125; ,19].

#### Eye-tracking data

1) *Pupil-based single eye data cleaning*: gaze samples out of the screen were excluded and each eye’s standard deviation (std) was computed for each trial and trial phase, and the ratio between the two eyes’ std was computed to determine if an eye’s data should be excluded (ratio of 1.82 or more, or 75% samples or more missing). The whole segment was excluded if ≥50% samples were missing for both eyes. 2) *Overall data cleaning*: wherever gaps in the data were found, 15 samples (30ms) on each side of the gap was removed to avoid boundary effects, and gaps were linearly interpolated if they were 160ms or shorter and the x and y coordinates at the start and end of the gap were less than 2% apart with respect to the overall screen size. The data was then smoothed with a moving average of 20 samples (40ms), and the gaze position was averaged over both eyes (where available). 3) *Parsing of fixations and saccade*s (*Exploration phase*): a velocity-based two-threshold method [20] was used to parse fixations and saccades for x and y coordinates of gaze location separately, using the sum of absolute changes between consecutive samples as the measure of velocity. Samples exceeding a high-threshold of 0.5 times the velocity’s std were identified as saccades and extended until the velocity fell below a low-threshold of 0.3. Periods between saccades lasting for at least 100ms and with less than 50% missing data were identified as fixations, and the fixations’ location was marked as belonging to one of the AOI for the visible cards around each card or outside AOIs. Only fixations within up-face AOIs during the *exploration phase* counted as exploratory fixations. 4) *Parsing of comparative looks (Exploration phase)*: any (direct or indirect) transition from a fixation in the AOI of one face-up card to a fixation in the AOI of the other face-up card during the was counted as a comparative look, and the first entry point in one of the two face-up AOIs counted as the first look. 5) *Calculation of decision slowing (Decision phase)*: decision slowing was operationalized as the duration between the time of the top card’s image reveal and the decision time (beginning of the fixation that triggered the selected card’s reveal). 6) *Within-group condition comparison:* trials following a correct choice were compared to trials following an incorrect choice using Bayesian statistics, testing for post-error increases i.e., enhanced exploration and decision slowing (unidirectional hypothesis: *After Incorrect > After Correct*).

#### EEG data

*1) Data cleaning and pre-processing in NetStation:* participants’ continuous EEG signal was filtered between 0.5Hz and 30Hz with a notch filter at 50Hz, segmented into epochs for ERN (-300ms to +550ms, aligned to the beginning of the fixation that triggered the selected card’s reveal) and FRN (-1,000ms to + 1,000ms aligned to the selected card’s reveal), and visually inspected for artefacts. Segments containing more than ∼33% noisy channels or segments for which the ERN/FRN had been excluded were excluded from analyses. Noisy channels in kept segments were interpolated using the neighboring channels, and the data was re-referenced to the average using PARE correction [21]. *2) Pre-processing in Matlab using FieldTrip toolbox*: segments were baselined trial-by-trial using the trial’s average over the -300ms to -150ms ERN period (for both ERN and FRN segments), averaged per subject and condition (correct or incorrect) and over fronto-central electrodes (using the same group of electrodes as identified in [16]: F2, F2-FCz, Fp2-Fz, Fz, Fz-FC1, Fz-Fp1, F1). *3) Whole-group cluster-based permutation analysis in Matlab with FieldTrip*: time-windows in which signal following *incorrect choices* was significantly more negative than following *correct choices* (directional one-tailed hypothesis) were identified within the whole group using cluster-based permutation analysis using a t-sum metric for N = 5,000 permutations and alpha=0.05. *4) Within-group condition comparison:* the peak (minimum) values per individual and condition within the time-windows of interest (identified by the cluster permutation tests) were extracted, and conditions were compared using Bayesian statistics testing for a post-error negativity effect (unidirectional hypothesis: *Correct > Incorrect*).

#### Performance data

Due to the adaptive difficulty design, infants’ accuracy was artificially maintained around 50%. Thus, to probe infants’ performance, a measure of their learning curve was used, calculated as the average local derivative in infants’ overall accuracy score (number of correct minus incorrect trials over trial number). Accuracy (the proportion of correct out of the total number of trials) is still reported for reference in the supplementary material’s descriptives.

#### Demographics data

Infants age and their general development were recorded as control measures. The Ages and Stages Questionnaire Third Edition [ASQ-3; 22] was used to measure general development at both visits using the 12- and 18-month versions. Questionnaires were completed by the caretakers online within a few days of the visit.

#### Inclusion and exclusion criteria

1) *Eye-tracking*: trials were excluded if the difficulty level changed between the preceding trial and the current one, if either the *exploration* or *decision* phase was rejected based on missing data, if the experimenter intervened (for example by introducing a break video), or if the trial represented a statistical outlier within a participant value for exploratory fixations or decision slowing over 6 Median Absolute Deviations or comparative looks value with a frequency lower than 5%). Furthermore, participants were excluded if they did not contribute any EEG data or if they were a statistical outlier within their *S* or *NS* group (value for exploratory fixations or decision slowing over 6 Median Absolute Deviations [same as trial rejection] or comparative looks values falling within the population’s 1^st^ or last percentiles); no participant was found to be a statistical outlier with this method, 58 participants were included (same as EEG). 2) *EEG*: trials were excluded based on noise level (bad trials marked by visual inspection of the ERN and FRN segments separately; only trials for which both the ERN and the FRN segment were marked as good were included), and participants contributing less than 6 included trials per condition (correct, incorrect) were excluded. This resulted in 58 participants included in the analysis for whom there was an average of 9.77 included trials per condition (std = 3.437 min. = 6, max. = 22 trials) and 3.09 excluded trials per condition (std = 2.31, min. = 0, max. = 13 trials), which did not significantly differ by group for *S* and *NS* participants (respectively, p=0.893 and p=0.100). Moreover, 67 participants were excluded: 11 due to technical issues, and 56 due to contributing too few or too noisy data, mainly due to fussiness related to wearing the EEG net.

## Results

### 12-month-olds internally monitor own errors

At their first visit, 12-month-old infants took part in a novel fully infant-controlled, decision-making task, based on match-to-sample paradigms [23,24]. Infants interacted with a screen via an eye-tracker, which used gaze position to trigger parts of the display in response to infants’ eye movements, in real time. This gave infants control of the task’s timing and progression and enabled free exploration and decision-making. In each test trial, infants were presented with 3 cards, 2 of which were initially facing up, and 1 facing down (Figure 1). The face-up cards each displayed a different image of items familiar to infants, and infants could look at them for up to 20s (*exploration phase*). Once infants gazed at the third (face-down) card, their gaze triggered the reveal of the card’s image, while simultaneously hiding the former two. The image on the third card always matched the image on one of the initially seen cards (now facing down) and was presented for up to 10s (*decision phase*). During this time, infants could gaze back at either of the former two cards and reveal their image again (*feedback phase*), thereby making a match (correct choice) or not (incorrect choice). Crucially, for infants to reveal the image of a selected card, they had to gaze at it for a cumulative 500ms. This allowed us to extract infants’ neural activity in the period after they had made their decision (their gaze landed on a face-down card) but before they had received any external feedback on their choice (image revealed).

To investigate internal monitoring of errors, we examined error-related negativity (ERN), a comparison of frontocentral neural activity following correct and incorrect decisions. If infants in the current study understood the match-to-sample rule, and internally monitored own decisions, the neural activity, aligned to the moment infants’ gaze landed on the selected card, was expected to be more negative following incorrect as compared to correct choices (directional one-tailed hypothesis). Data from *N*=58 infants (see detailed descriptive values in Supplementary Materials) indeed revealed an ERN (Figure 2): cluster-based permutation analysis revealed two time periods in which infants’ neural responses to incorrect choices were significantly lower than responses to correct choices, an early peak at 33-172ms post-choice (*p*=0.031), and a late peak at 332-547ms (*p*=0.013). Both peaks matched the ERN previously observed in infants [16], while only the early peak matched the timing and polarity of the adult ERN, thus the first peak was retained for the remainder of the analyses. To further validate that infants could distinguish between correct and incorrect choices, we additionally analysed infants’ neural responses to external feedback (aligned to the reveal of the selected card’s image). The same procedure revealed a feedback related negativity effect (FRN) over the same scalp region (two time periods were identified, early peak: 46-298ms, *p*=0.017; later peak: 457-930ms, *p*=0.001, the latter of which matched the timing of the adult FRN and was thus retained for further analyses). Combined, the observed neural responses suggest that 12-month-olds distinguished between correct and incorrect choices they had made, based both on external feedback (FRN) and, crucially, even prior to receiving the feedback (ERN).

**Figure 2.**
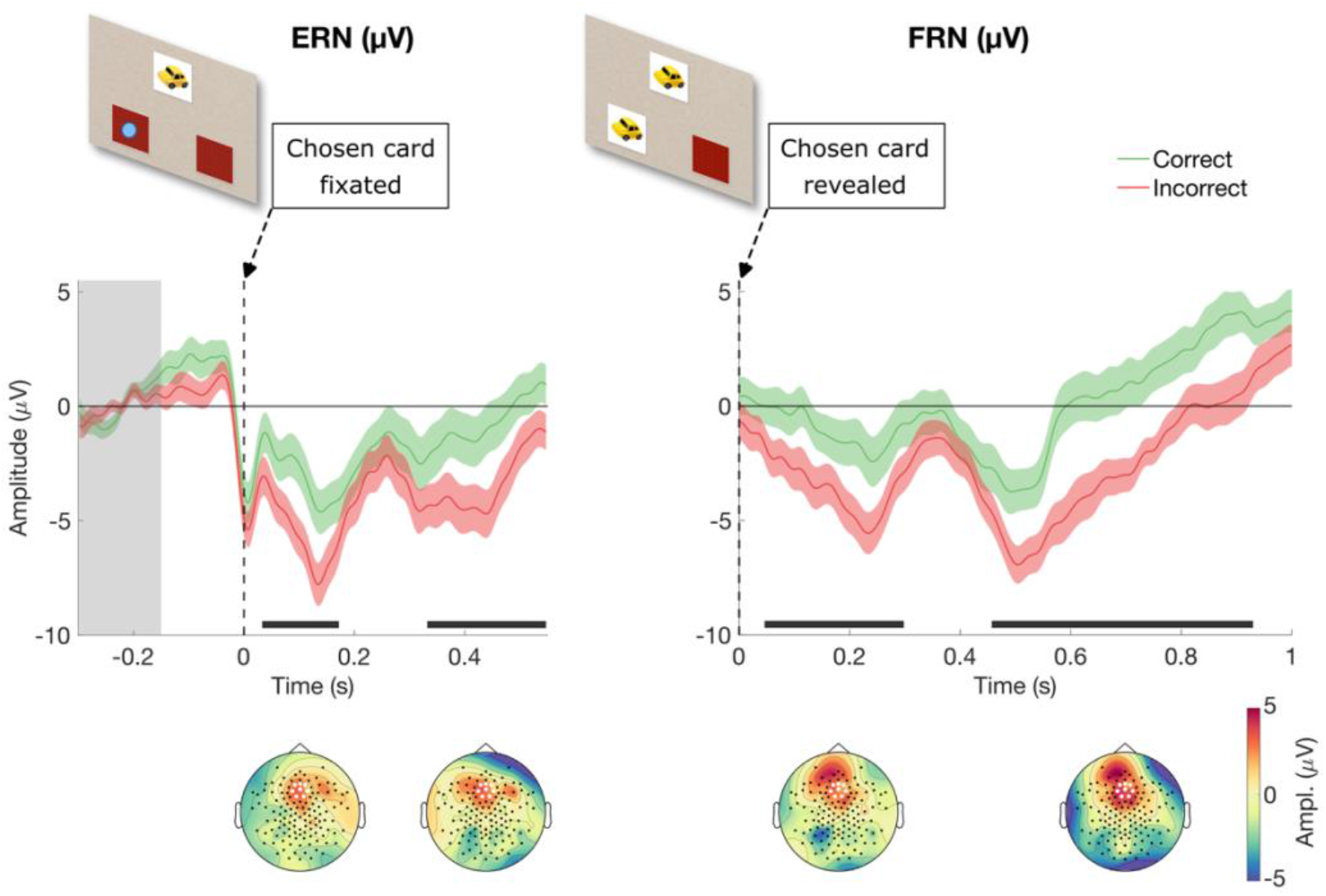
ERN and FRN responses. Average ERPs are represented with lines, with shading representing the SEM. Data was aligned to the moment a chosen card was fixated (ERN, left), or the moment the chosen card was revealed (FRN, right), as depicted on top (the blue dot signifies the infant gaze). Data was baselined from -300 to -50ms (shaded area). Solid lines at the bottom of the plots represent the time windows identified as significant by the time-cluster analysis. Topographies of the signal within the time window of interest are depicted at the bottom.

### Error monitoring predicts mirror self-recognition

Having established that 12-month-olds exhibit internal error-monitoring in our paradigm, we investigated whether this capacity predicts the later emergence of self-representation, by testing the same infants 6 months later, using the mirror self-recognition (MSR) task. We used a binary score on the MSR as the dependent outcome (success: *S* group, *N*=33; or not: *NS* group, *N*=20), and the magnitude of the ERN as the predictor (peak amplitude of frontocentral potentials for correct minus incorrect choices). A significant logistic regression model emerged (*p*=0.031): ERN at 12 months was found to correctly predict infants’ self-representation 6 months later with 69.8% accuracy, correctly classifying 90.9% of *S* and 35.0% *NS* infants. Importantly, while the target behaviour in the MSR task (touching the mark on own face) is taken as evidence of self-representation, the absence of this behaviour is inconclusive. The *NS* group is therefore expected to be more heterogenous and thus be classified with lower accuracy. To account for possible differences in infants’ general development, a further stepwise model was conducted, with infants’ age and a standardized measure of cognitive development (ASQ) as additional predictors. Both variables were rejected as relevant predictors of MSR, and anecdotal evidence was found for no group differences in either measure (age: BF01=2.82; general development: BF01=2.55). These results thus indicate that the more sensitivity an infant show in internally monitoring their own mistakes at 12 months (the stronger the ERN response), the more likely they are to have acquired a conceptual self-representation by the age of 18 months, irrespective of their age or cognitive development. Finally, no evidence was found for a predictive effect of the FRN on later self-representation (p=0.946), suggesting it is specifically infants’ internal (and not external) error-monitoring that is predictive of the later emergence of self-representation.

To further characterize the developmental relationship found, we analyse infants’ internal monitoring capacities separately for infants that showed self-representation at 18 months (S group) and those that did not (NS group; Figure 3). Peak analysis of the ERN response (37-172ms) revealed very strong evidence of internal error-monitoring (directional one-tailed hypothesis) in the S group only (BF10=230.33). In contrast, moderate evidence for the absence of an ERN was found in the NS group (BF01=4.19). Importantly, this cannot be explained by differences in data quality or group size, as moderate evidence for the FRN effect was found in both groups (S: BF10=4.22; NS: BF10=6.15) with moderate evidence for no group difference (BF01=3.53). These results thus indicate that both groups of infants distinguished between correct and incorrect choices based on feedback, but only the S group additionally monitored their own decisions internally.

**Figure 3.**
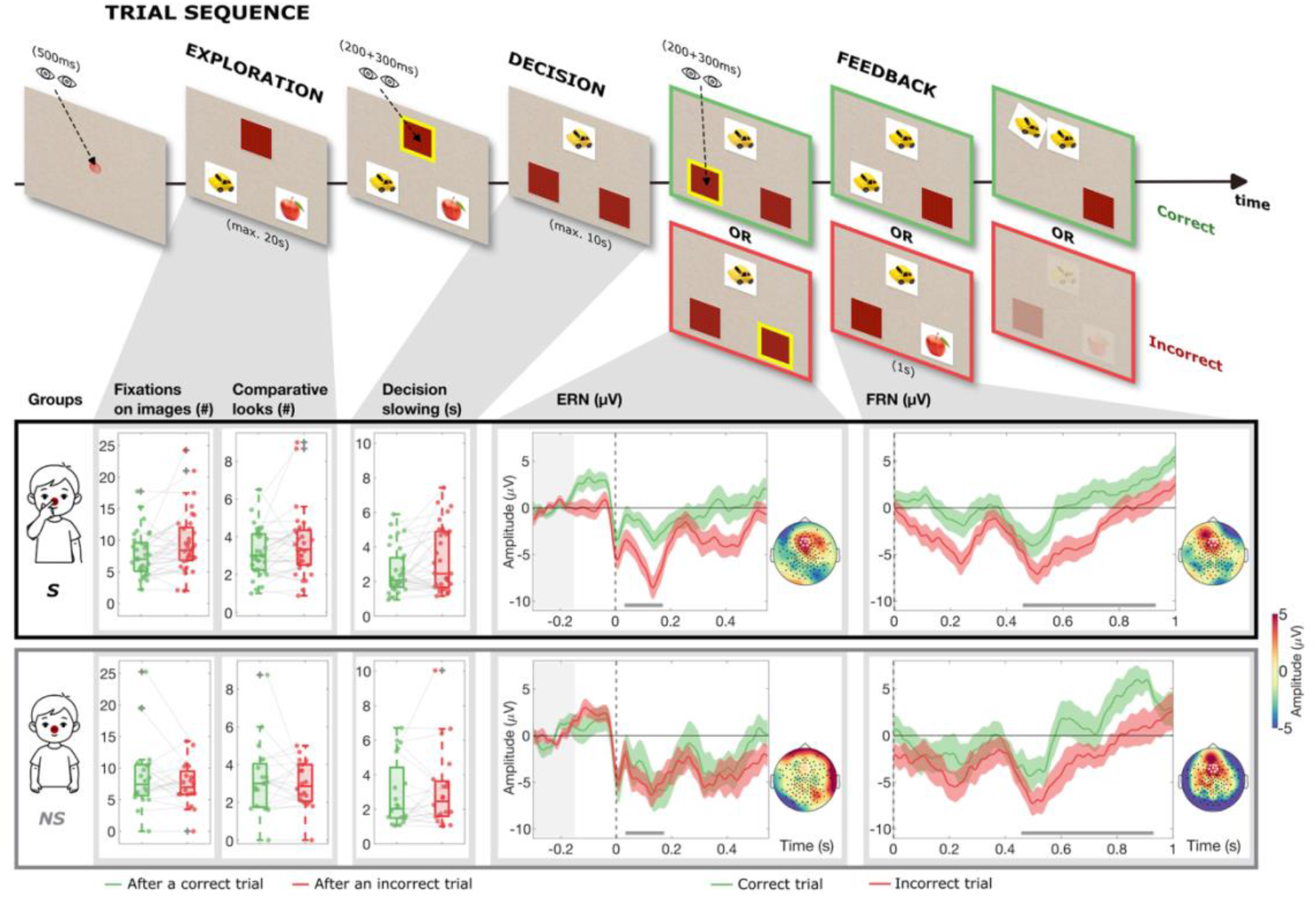
Task design and results split by groups. Trial sequence (top): images of the different trial steps are depicted along the 3 phases (exploration, decision, feedback) with eye symbols representing the gaze-contingent parts of the trials. Green and red highlights around images represent the sequence for correct or incorrect choices respectively. Results for the different eye-tracking and EEG measures at 12 months are presented under the trial phase at which they were extracted, for the two MSR groups separately: the group who showed evidence of self-representation at 18 months (*S*) in the black box; and the group who did not (*NS*) in the grey box. Correct (or following correct) trials are depicted in green, while incorrect (or following incorrect) trials are depicted in red. Eye tracking data (left) is presented in boxplots: boxes represent 25 to 75 percentiles, lines depict the median, and whiskers represent the range of included data points. Individual data points are represented by dots, and outliers are depicted by crosses. Average ERPs (right) are represented with lines, with shading representing the SEM. Data was baselined from -300 to -50ms (shaded area). Solid lines at the bottom of the plots represent the time windows identified as significant by the initial time-cluster analysis, while topographies of the signal within the time window of interest for each group and signal (ERN or FRN) are plotted to the right of each plot.

### Error monitoring guides adjustments in information-seeking and decision-making

Finally, we explored whether internal error-monitoring not only predicts infants’ self-representation, but also associates with concurrent behavioural adjustments on task. Recorded as part of the same experiment at 12 moths of age, infants’ eyetracking data was analysed for evidence of two types of post-error adjustments. First, the *decision phase* was analysed for evidence of post-error slowing, the commonly observed effect of increased decision times following the commitment of an error [25,26]. Second, task-specific adjustments were investigated in the *exploration phase,* namely increased post-error information gathering in the form of fixations on, and comparative looks between, the displayed images. Comparing trials following an error to trials following a successful match (Figure 3), we found that only infants who evidenced internal error-monitoring and later self-representation (*S* group) showed evidence of behavioural adjustments: moderate evidence of post-error decision slowing (BF10=5.11), and strong evidence of increased post-error fixations (BF10=10.47), but no conclusive evidence of increased post-error comparative looks (BF10=1.40). In contrast, the *NS* group evidenced: moderate evidence for the absence of an increase in fixations and comparative looks following errors (BF01=9.11 and BF01=6.84, respectively), and anecdotal evidence against a decision slowing (BF01=2.38). Furthermore, in the *S* group, moderate evidence was found for a positive relationship between infants’ ERN magnitude and their performance on task (average local derivative of infants’ overall accuracy score, Figure 4), suggesting that the capacity to internally monitor own errors played a functional role in infants’ overall performance on the task (R=0.420, BF10=6.70). In contrast, moderate evidence for the absence of a relationship between ERN and performance was found in the *NS* group (R=-0.229, BF01=6.55). Interestingly, the absence of ERN and behavioural adjustments in the *NS* group was not found to result in worse performance overall (anecdotal evidence for no group difference, BF01=2.38). Comparable performance may thus be achieved via routes other than the internal error monitoring and behavioural adjustments probed in this study. Combined, our task’s neural and behavioural measures provide converging evidence suggesting that, already at 12 months of age, infants’ internal error-monitoring can aid performance and can be utilized for post-error behavioural adaptations. Crucially, this was only the case for infants who also exhibited evidence of self-awareness at 18 months.

**Figure 4.**
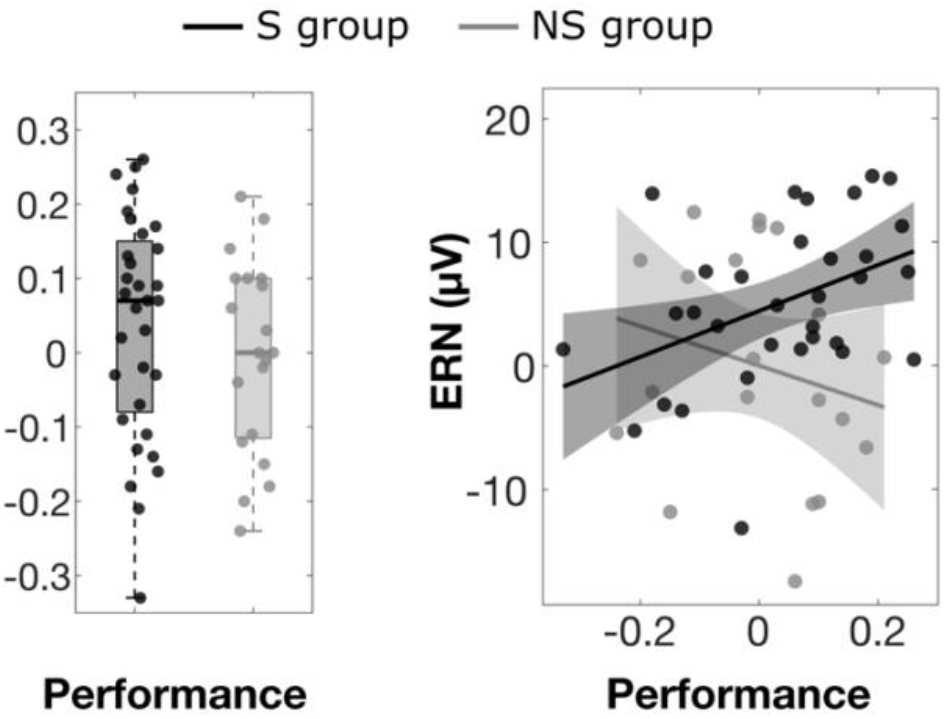
Performance and ERN. The groups’ performance (average local slope in the participant’s difficulty level as a function of trial number) is depicted for the *S* group in black and the *NS* group in grey (left), together with the groups’ correlation between ERN amplitude and performance (right).

## Discussion

Our findings revealed that preverbal human infants’ capacity to internally monitor their own errors developmentally predicts the emergence of their conceptual self-representation 6 months later. This relationship was specific to internal error-monitoring, and not external feedback processing, and could not be explained by infants’ age or general cognitive development. Moreover, only the group of infants, who evidenced conceptual self-representation at 18 months, also demonstrated a clear ERN as well as early behavioural adjustments following errors at 12 months. Combined, these converging neural and behavioural results indicate that error-detection, if available, can be utilized for adaptive information-seeking and decision-making as early as 12 months of age. Finally, the magnitude of infants’ neural response to errors correlated with their performance on task, but did not result in significantly different overall performance, when compared to infants who did not evidence internal error-monitoring nor later self-representation. This suggests that the relationship found between early error-monitoring and later conceptual self-representation is unlikely a reflection of differences in infants’ general cognitive abilities but appears specific to infants’ developing capacities of self-reflection.

Given that, in adults, neural signals of error detection can be recorded in absence of a subjective report of having made an error [27], it is unclear whether error detection in infants occurs on a conscious level [16,28]. Consequently, the observed adjustments in infants’ behaviour, namely increased information seeking and slowed decision making, may reflect exploratory or corrective behaviours that are directly triggered by first order neural representations. According to the proposal of a ‘model free’ metacognition, one can postulate neural representations which are metacognitive in the sense that they represent uncertainty or correctness conditions but do not necessitate any kind of model of one’s own mind [7,8]. Thus, infants’ (and other non-human animals’) information seeking behaviour may be appropriately guided in absence of self-awareness and of an additional, second-order meta-representation of own errors or uncertainty. However, the fact that the magnitude of infants’ response to own errors at 12 months predicted which of the infants will have evidenced self-representation 6 months later, strongly suggests that, at least in humans, these processes are not independent. Moreover, it seems logically necessary that error-detection signals must (eventually) be consciously accessed, if error-monitoring is to be utilised for, or integrated into, a conceptual self-representation. One challenge of specifying the process and mechanisms of building a conceptual self-representation thus lies in specifying the mechanism of conscious access to mental representations itself. Both adult as well as developmental theories have put forward proposals as to how the mind may generate conscious experience [29], or turn information that is represented in the mind into progressively more explicit information that is accessible to the mind [30], however, empirical evidence is yet to provide conclusive data regarding the mechanisms supporting such transformations.

One possibility for how error-monitoring may contribute to the building of a conceptual self-representation is that early availability of internal error signals can lead to better tuned metacognitive feelings. Defined as affective responses stemming from monitoring the successes and failures of own cognitive actions [31], metacognitive feelings can initiate epistemic behaviours, such as targeted information sampling. Such error-guided information seeking could facilitate the gradual building of conceptual representations in general, including the concept of self specifically. Alternatively, but similarly grounded in error-guided information seeking, proposals emphasising one’s tracking of own learning progress as the mechanism determining which activities an individual will be motivated to pursue [32] would propose that information related to, and generated by, the self is particularly informative and allows for comparatively rapid learning progress. This can thereby drive self-related information seeking, and consequently the early emergence of a category of self [33]. Although learning progress account does not distinguish between internally generated error signals and external feedback, it is plausible that the capacity of detecting errors as one’s own would facilitate meta-learning, that is making use of one’s prior experiences to learn how to learn better in the future [34]. The behavioural adjustments following errors, seen in infants who evidenced internal error monitoring in our study, may therefore reflect, or ultimately lead to, infants meta-learning about their own information-seeking and decision-making strategies. As such, error detection and internal evaluations of own learning may themselves become an integral part of the conceptual understanding of the self as an epistemic agent, as previously hinted at by adult research [35,36].

Finally, as the current evidence demonstrates, error-detection can lead to appropriate online adjustments in information-seeking and decision-making and can facilitate task performance even prior to any evidence of a conceptual self-representation. This raises another open question, namely, if cognitive control and adaptive learning can operate in absence of a conceptual self, why would humans nonetheless develop a conceptual self-representation or explicit forms of metacognition? In addressing this question, some theoretical proposals have focused on plausible evolutionary pressures for evolving explicit reflective capacities and proposed the human-specific needs to reason about others’ minds [37] and to coordinate mental content between multiple individuals [38] as the best candidates. For example, it is argued that the capacity to share our minds with others is instrumental in creating and sustaining the complex social structures and cumulative culture characteristic of our species [38]. Other, mainly developmental, theories that have focused on the human-specific input that may trigger a conceptual self to develop have likewise emphasised social interactions, identifying direct eye gaze [14] and turn-taking contingencies [39] as plausibly involved in the gradual emergence of a conceptual self. Interestingly, empirical evidence from children suggests that individuals’ response to errors (as reflected in the magnitude of the ERN) may similarly be modulated by different aspects of social contexts [40]. This raises the intriguing new possibility that one way in which early social input may affect the development of a self-representation is through modulating infants’ detection and sensitivity to own errors.

This study revealed a developmental bridge between error-monitoring and self-representation in human infants, providing initial empirical traction on questions that have fascinated philosophers and cognitive scientists for centuries: whether metacognition implies self-awareness, is independent of it, or whether one emerges as a consequence of the other. The developmentally predictive link recorded here supports the latter option, whereby metacognitive processes likely contribute to the building of a later emerging conceptual representation of oneself as a distinct agent.

**Transparency statements**

## Acknowledgements

We thank our funders and all the participating families, students, and colleagues who made this research possible.

## Ethics

This study was approved by the Ethical Committee of the University of Copenhagen’s Institute of Psychology (Approval No. IP-EC-1512-2-2022).

## Preregistration & Preprint

The aims, hypotheses, methods, variables and analyses were preregistered on OSF before the data was accessed (https://osf.io/gcb5z/overview?view_only=b37adcf37c664517be43685c40a3c5c2 ), with deviations described in Supplementary Table S5. Early versions of this paper were preprinted on BioRxiv (https://doi.org/10.64898/2026.02.01.702903).

## Data & Code Availability

Access to the primary data is restricted; requests to the corresponding author can be made, pending agreements for sharing of sensitive (infant) data. All experimental and analysis scripts are publicly available: (https://github.com/earlychildcog/curiosity-error-stimpres https://github.com/earlychildcog/curiosity-error-analysis).

## Authors Contribution

Conceptualization: C & K; Data Curation: C & D; Formal Analysis: C & D; Funding Acquisition: K; Investigation: C; Methodology: C & D, K; Project Administration: C; Resources: K; Software: C & D; Supervision: C & K; Validation: K, C & D; Visualization: C; Writing – Original Draft: C & K; Writing – Review & Editing: C, D & K.

