## Supplementary materials for "I err, therefore I am: Neural signature of error-monitoring predicts self-awareness in infants"

### Descriptive statistics

Table S1. Whole-group EEG descriptive statistics.

| Variable | Trial set | Group | Mean | Std. | Min. | Max. |
|---|---|---|---|---|---|---|
| **ERN** (uV) | **Correct** | **all** | -7.14 | 6.78 | -20.43 | 6.69 |
| | **Incorrect** | **all** | -10.20 | 6.53 | -28.87 | 1.67 |
| | **Difference** | **all** | 3.06 | 8.30 | -17.42 | 20.51 |
| **FRN** (uV) | **Correct** | **all** | -8.64 | 6.50 | -21.26 | 4.64 |
| | **Incorrect** | **all** | -11.87 | 5.61 | -25.97 | 3.28 |
| | **Difference** | **all** | 3.22 | 7.22 | -8.46 | 24.95 |

Table S2. Whole-group additional EEG descriptive statistics (provided for reference on additional time-windows identified in the cluster analysis but not retained for the main analysis).

| Variable | Trial set | Group | Mean | Std. | Min. | Max. |
|---|---|---|---|---|---|---|
| *2nd window ERN* (uV) | **Correct** | **all** | -6.31 | 8.34 | -27.12 | 17.23 |
| | **Incorrect** | **all** | -9.27 | 7.02 | -29.79 | 2.95 |
| | **Difference** | **all** | 2.96 | 9.35 | -17.31 | 36.95 |
| *1st window FRN* (uV) | **Correct** | **all** | -7.23 | 6.54 | 26.69 | 8.90 |
| | **Incorrect** | **all** | -9.63 | 6.16 | -22.92 | 6.19 |
| | **Difference** | **all** | 2.39 | 8.33 | -16.28 | 22.61 |

Table S3. Per-group EEG & eye-tracking descriptive and Bayesian statistics for within-group comparisons (unidirectional: H1 is defined as *Correct* > *Incorrect* or *After Incorrect* > *After Correct*); significant comparisons (BF > 3) are depicted in bold.

| Variable | Group | Trial set | Mean | Std | Min. | Max. | $BF_{10}$ | $BF_{01}$ |
|---|---|---|---|---|---|---|---|---|
| **ERN** (uV) | ***NS*** | **Correct** | -8.86 | 8.50 | -20.43 | 5.05 | 0.240 | **4.19** |
| | | **Incorrect** | -8.93 | 5.68 | -19.97 | 1.52 | | |
| | | **Difference** | 0.07 | 9.06 | -17.42 | 12.44 | - | - |
| | ***S*** | **Correct** | -5.89 | 5.67 | -18.73 | 6.69 | **230.33** | 0.01 |
| | | **Incorrect** | -10.72 | 6.47 | -24.49 | 1.67 | | |
| | | **Difference** | 4.84 | 6.72 | -13.11 | 15.38 | - | - |
| **FRN** (uV) | ***NS*** | **Correct** | -8.82 | 7.07 | -21.26 | 2.44 | **6.15** | 0.16 |
| | | **Incorrect** | -12.37 | 5.01 | -25.35 | -1.92 | | |
| | | **Difference** | 3.54 | 6.14 | -8.46 | 12.96 | - | - |
| | ***S*** | **Correct** | -8.41 | 6.41 | -19.05 | 4.64 | **4.22** | 0.24 |
| | | **Incorrect** | -11.81 | 6.31 | -25.97 | 3.29 | | |
| | | **Difference** | 3.40 | 8.21 | -8.06 | 24.95 | - | - |
| **Exploratory Fixations** (on face-up cards during the exploration phase) | ***NS*** | **After Correct** | 8.64 | 5.55 | 0.00 | 25.25 | 0.11 | **9.11** |
| | | **After Incorrect** | 7.34 | 3.45 | 0.00 | 14.25 | | |
| | | **Difference** | -1.30 | 4.27 | -11.58 | 4.25 | - | - |
| | ***S*** | **After Correct** | 7.72 | 3.54 | 2.20 | 17.75 | **10.47** | 0.10 |
| | | **After Incorrect** | 9.59 | 4.89 | 2.00 | 24.25 | | |
| | | **Difference** | 1.86 | 3.78 | -5.90 | 13.00 | - | - |
| **Comparative Looks** (between face-up cards during the exploration phase) | ***NS*** | **After Correct** | 3.22 | 1.88 | 0.00 | 8.75 | 0.15 | **6.85** |
| | | **After Incorrect** | 2.97 | 1.22 | 0.00 | 5.00 | | |
| | | **Difference** | -0.25 | 1.55 | -4.75 | 2.58 | - | - |
| | ***S*** | **After Correct** | 3.15 | 1.25 | 1.00 | 6.50 | 1.40 | 0.72 |
| | | **After Incorrect** | 3.57 | 1.75 | 0.87 | 9.00 | | |
| | | **Difference** | 0.42 | 1.38 | -3.25 | 4.50 | - | - |
| **Decision Slowing** (s) | ***NS*** | **After Correct** | 2868.75 | 1917.86 | 1058.40 | 6712.33 | 0.42 | 2.38 |
| | | **After Incorrect** | 3122.61 | 2299.56 | 1016.43 | 10038.00 | | |
| | | **Difference** | 253.87 | 1674.93 | -2661.67 | 4522.00 | - | - |
| | ***S*** | **After Correct** | 2581.54 | 1329.38 | 941.00 | 5887.60 | **5.11** | 0.20 |
| | | **After Incorrect** | 3255.27 | 1857.61 | 1156.40 | 7428.67 | | |
| | | **Difference** | 673.73 | 1561.21 | -2674.60 | 4167.80 | - | - |

Table S4. Per-group performance & demographics descriptive and Bayesian statistics for between-group comparisons (bidirectional: H1 is defined as *S* > *NS* or *NS* > *S*); significant comparisons (BF > 3) are depicted in bold.

| Variable | Group | Mean | Std | Min. | Max. | $BF_{10}$ | $BF_{01}$ |
|---|---|---|---|---|---|---|---|
| **Performance** | ***NS*** | -0.01 | 0.13 | -0.24 | 0.21 | 0.42 | 2.380 |
| | ***S*** | 0.04 | 0.15 | -0.33 | 0.26 | | |
| **Accuracy** | ***NS*** | 0.50 | 0.08 | 0.36 | 0.64 | 0.35 | 2.67 |
| | ***S*** | 0.52 | 0.08 | 0.33 | 0.656 | | |
| **Number of test trials** | ***NS*** | 36.15 | 8.90 | 23.00 | 58.00 | 0.34 | 2.90 |
| | ***S*** | 34.58 | 7.40 | 22.00 | 51.00 | | |
| **Number of Easy trials** | ***NS*** | 15.75 | 9.24 | 2.00 | 31.00 | 0.31 | 2.96 |
| | ***S*** | 14.5 | 10.40 | 3.00 | 37.00 | | |
| **Number of Medium trials** | ***NS*** | 11.95 | 6.10 | 2.00 | 29.00 | 0.38 | 2.87 |
| | ***S*** | 10.54 | 5.92 | 2.00 | 27.00 | | |
| **Number of Hard trials** | ***NS*** | 6.15 | 6.61 | 0.00 | 19.00 | 0.30 | **3.46** |
| | ***S*** | 5.61 | 6.22 | 0.00 | 22.00 | | |
| **Number of Extra-Hard trials** | ***NS*** | 2.30 | 5.19 | 0.00 | 16.00 | 0.41 | 2.33 |
| | ***S*** | 3.97 | 6.80 | 0.00 | 24.00 | | |
| **Age** (days) | ***NS*** | 369.70 | 7.15 | 351.00 | 378.00 | 0.40 | 2.82 |
| | ***S*** | 367.36 | 9.67 | 349.00 | 384.00 | | |
| **Developmental stage** (total ASQ score) | ***NS*** | 205.25 | 36.15 | 125.00 | 260.00 | 0.39 | 2.55 |
| | ***S*** | 215.30 | 43.20 | 140.00 | 295.00 | | |

Table S5: Preregistration deviations (adapted from Willroth & Atherton, 2023).

| Details | | Original wording | Deviation description | Extent of deviation | Judgement of impact |
|---|---|---|---|---|---|
| Type | Analyses | One main objective was preregistered:<br>"We investigate whether 12-month-old infants adapt their information gathering in relation to different learning contexts, and how infants' information seeking relates to their processing of their own errors."<br>With three main questions:<br>"Do infants adapt their information search (visual scanning) depending on the difficulty level (similarity of images)?"<br>"Do infants detect that they have selected a mismatching card even before they receive visual feedback (neural responses)?"<br>"How do(es) infants' (internal monitoring of own) errors relate to their information search?" | Due to low trial numbers in higher difficulty levels and due to the adaptive difficulty design making the variable tightly dependent on infant's behaviour and thus rather circular, the Question 1 could not be examined, while Questions 2 and 3 remain central to the present paper.<br>All our eye-tracking and ERN variables were preregistered.<br>Several other measures were listed in the preregistration as potential variables of interest but were not pursued due to data characteristics - a possibility which was acknowledged as likely in the preregistration due to the novelty of the paradigm and of the collected measures (Preregistration, Section 2, final paragraph).<br>The self-recognition score was listed as a potential additional variable of interest (Preregistration, Section 8, "self-recognition score on the rouge test") prior being collected at the second visit of the longitudinal study, but became a core interest of the current manuscript. | Major & Minor | Minor:<br>Not exploring Question 1 (difficulty-related adaptation) due to practical impossibilities and only exploring Questions 2 and 3 (error-detection and error-related adaptation of exploration) was sufficient to explore the main objective of how infants adapt their information gathering based on their own errors. The main objective remained attainable. All variables used were pre-registered. This is considered a minor deviation.<br>Major:<br>One of the potential additional objectives preregistered (relation to self-recognition) took on a more central part in the current paper, which is considered a major (although preregistered) deviation. |
| Reason | Plan not possible | | | | |
| Timing | After data access | | | | |
